# Comprehensive longitudinal study of epigenetic mutations in aging

**DOI:** 10.1101/744250

**Authors:** Yunzhang Wang, Robert Karlsson, Juulia Jylhävä, Åsa K. Hedman, Catarina Almqvist, Ida K. Karlsson, Nancy L. Pedersen, Malin Almgren, Sara Hägg

## Abstract

**Background:** The role of DNA methylation in aging has been widely studied. However, epigenetic mutations, here defined as aberrant methylation levels compared to the distribution in a population, are less understood. Hence, we investigated longitudinal accumulation of epigenetic mutations, using 994 blood samples collected at up to five time points from 375 individuals in old ages.

**Results:** We verified earlier cross-sectional evidence on the increase of epigenetic mutations with age, and identified important contributing factors including sex, CD19+ B cells, genetic background, cancer diagnosis and technical artifacts. We further classified epigenetic mutations into High/Low Methylation Outliers (HMO/LMO) according to their changes in methylation, and specifically studied methylation sites (CpGs) that were prone to mutate (frequently mutated CpGs). We validated four epigenetically mutated CpGs using pyrosequencing in 93 samples. Furthermore, by using twins, we concluded that the age-related accumulation of epigenetic mutations was not related to genetic factors, hence driven by stochastic or environmental effects.

**Conclusions:** Here we conducted a comprehensive study of epigenetic mutation and highlighted its important role in aging process and cancer development.

## Introduction

Epigenetic processes, among which DNA methylation is one of the most well studied, are fundamental in human aging [1]. Studies on DNA methylation have identified age-associated changes in methylation levels shared by individuals [2,3], and have also reported an increasing divergence of methylation levels between individuals with age [4,5].

Epigenetic mutations, defined as aberrant methylation levels that can lead to unusual gene expression, may be involved in cancer development and important for human aging [6,7]. Unlike age-associated changes in methylation levels that are shared among individuals, the incidences of epigenetic mutations are rare, stochastic and inconsistent between individuals. Epigenetic mutations can partly explain the increasing variability of methylation levels between individuals over time, but the extreme methylation levels may concur stronger biological consequences, such as cancer. Epigenetic mutations could contribute to the aging process through the accumulation of abnormally methylated CpGs (cytosine-phosphatase-guanine sites), which could further cause abnormal gene expression and downstream effects in tissues. A previous study by Gentilini *et al* [7] specifically defined epigenetic mutations as extreme outliers within a population, with methylation levels exceeding three times interquartile ranges (IQR) of the first quartile (Q1-3 × IQR) or the third quartile (Q3+3 × IQR). They found that the total numbers of epigenetic mutations increased exponentially with age. However, since this finding was based on a cross-sectional study, it needs to be validated in a longitudinal setting, where the accumulation of epigenetic mutations over time can be followed within the same individuals. Moreover, it is not yet known what the clinical consequences of accumulated epigenetic mutations are, and if individuals with a high burden of epigenetic mutations are prone to develop cancer as previously suggested [6,8].

In this study, we used a Swedish twin cohort including 375 individuals sampled up to five times in late life across 18 years (Table 1). We first validated the age-related increase of epigenetic mutations from a longitudinal perspective. Next, we identified important factors associated with the number of epigenetic mutations, including sex, cellular composition (CD19 B-cells), genetic background and technical artifacts. In parallel, we analyzed the direction of change in methylation level and characterized the epigenetic mutations as High-(HMO) and Low Methylation Outliers (LMO). We also studied the association between epigenetic mutations and cancer, as well as the genetic influence on epigenetic mutations using a twin approach. Last, we validated a select set of epigenetic mutations using bisulfite pyrosequencing.

**Table 1.** Characteristics of study participants in SATSA (n=375 unique individuals).

| Longitudinal wave | Year of sample collection | Number of Participants (new recruits) | Female Proportion | Age mean (SD) |
| --- | --- | --- | --- | --- |
| 1 | 1992-1994 | 232 | 58% | 68.5 (9.1) |
| 2 | 1999-2001 | 239 (101) | 63% | 71.1 (10.1) |
| 3 | 2002-2004 | 186 (25) | 54% | 72.1 (9.1) |
| 4 | 2008-2010 | 183 (14) | 61% | 76.2 (8.5) |
| 5 | 2010-2012 | 154 (3) | 66% | 77.0 (8.4) |

SATSA: The Swedish Adoption/Twin Study of Aging

## Results

### Longitudinal accumulation of epigenetic mutations is exponentially associated with age

To explore the longitudinal increase in number of epigenetic mutations, we measured DNA methylation data (Illumina 450k array) repeatedly in whole blood samples (n=994) from participants in the Swedish Adoption/Twin Study of Aging (SATSA; Table 1) [9]. To avoid confounding by underlying genetic variation, we removed 20,660 CpGs that were associated with at least one single nucleotide polymorphism (SNP) (p<1e-14) within 1 Mbps (mega base pairs), i.e. cis-methylation quantitative loci (cis-meQTLs). In the remaining 370,234 CpGs, the number of epigenetic mutations ranged from 58 to 26,291 in each sample, using the definition in Gentilini *et al* [7]. Across samples, the number of epigenetic mutations had a right-skewed distribution, which was close to normal distribution after log10-transformation (Figure S1).

After identifying epigenetic mutations in SATSA, we found that the log10 total number of epigenetic mutations increased with age (p=1.22e-13) longitudinally (Figure 1A). We also identified additional factors and confounders associated with the number of epigenetic mutations (Table 2). Women had a slightly higher average number of epigenetic mutations than men (p=6.33e-3). Low sample quality, as defined by the log10-transformed number of CpGs with detection p-values over 0.01, was positively associated with the total number of epigenetic mutations (p=1.48e-117). In general, unreliable samples tended to have more epigenetic mutations, indicating that measurement errors could also be identified as epigenetic mutations. However, after adjusting the mixed models for detection p-value, the effect of age on number of epigenetic mutations remained unchanged. Using predicted cellular compositions, CD19+ B cell composition was positively associated with the total number of epigenetic mutations (p=5.06e-23). After removing cis-meQTLs, the first genetic principal component (PC) showed only a minor effect on the total number of epigenetic mutation (p=0.041).

**Figure 1.**
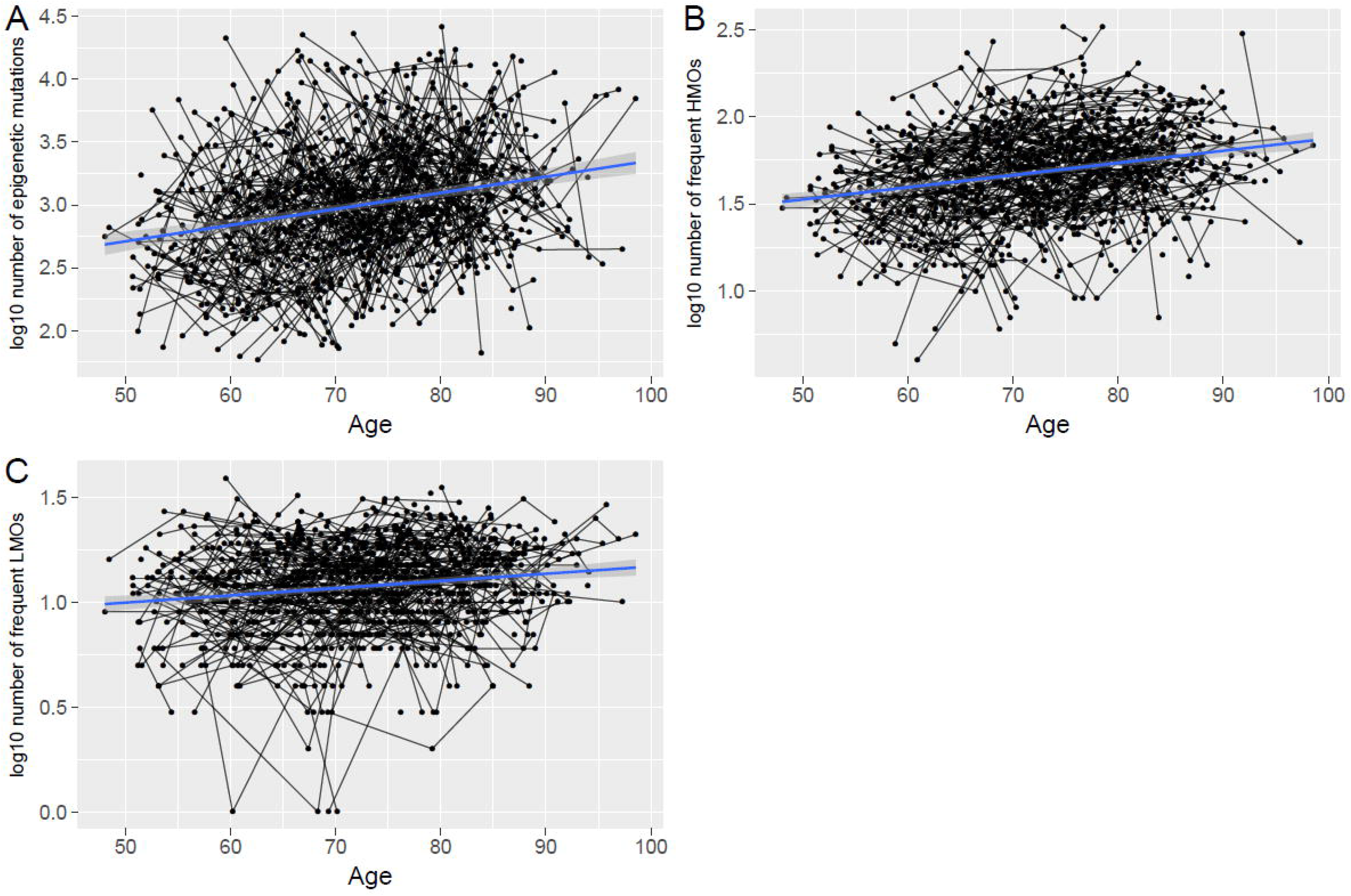
The number of epigenetic mutations (log10-transformed) increased longitudinally with age in a longitudinal perspective using genome-wide DNA methylation data from repeated whole blood samples collected in the Swedish Adoption/Twin Study of Aging (SATSA; n=375 participants). The numbers of epigenetic mutations of samples were counted from: A) total epigenetic mutations (n=370,234, p=1.22e-13 for association with age), B) frequent high methylation outliers (HMO) (n=969, p=2.09e-17 for association with age), and C) frequent low methylation outliers (LMO) (n=216, p=1.14e-05 for association with age).

Out of all CpGs, 237,398 (64%) were defined as epigenetic mutations in at least one sample, but only 1,185 (0.32%) CpGs were mutated in more than 50 samples; subsequently defined as frequently mutated CpGs. Only two of the 1,185 frequently mutated CpGs were also identified to be age-differentially methylated sites (aDMS) in our previous study [3]. The frequently mutated CpGs were still significantly associated with age, sample quality, CD19+ B cell compositions and genetic PC1, while sex was no longer significant (Table 2).

**Table 2.**
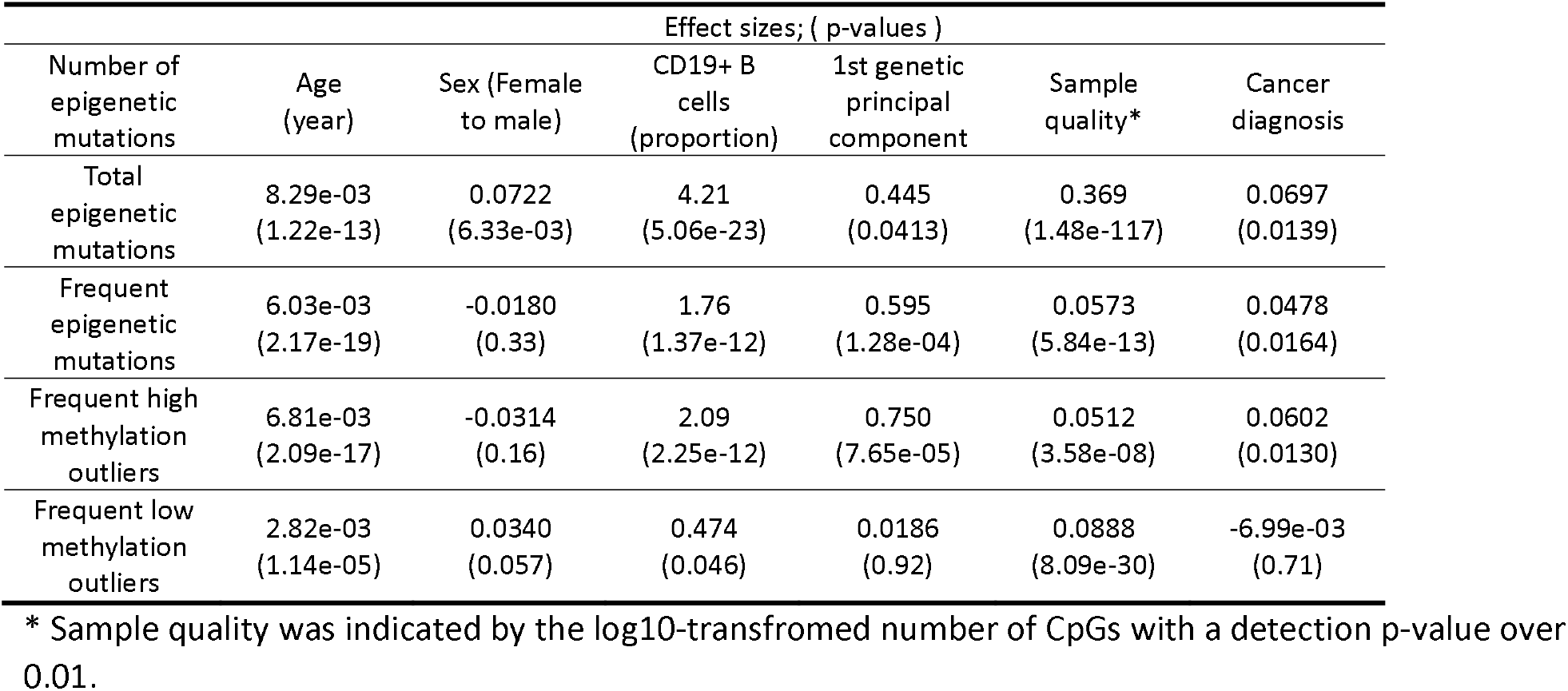
The association between number of epigenetic mutations (log10-transformed) and age from mixed models with confounders.

### High/Low Methylation Outliers

Compared to normal methylation levels in the population, epigenetic mutations can be either higher or lower in methylation level. Hence, we defined HMO and LMO as CpGs with abnormally higher or lower methylation levels than the average (Figure S2). Of the defined epigenetic mutation sites, almost half were identified as HMOs and the other half as LMOs (118,259 HMOs and 119,175 LMOs). Thirty-six CpGs were defined as both HMOs and LMOs because those sites had intermediate methylation levels and very small IQRs. However, among the frequently mutated CpGs, there were significantly more HMOs than LMOs (969 and 216, p<1e-16) (Figure 2). Nevertheless, numbers of both sets of frequent mutations (log10-transformed) significantly increased with age (p=2.09e-17 for HMOs and p=1.14e-05 for LMOs) (Figure 1B and C). Sex was no longer a significant factor with either frequent HMOs or LMOs. The composition of CD19+ B cell was still strongly associated with HMOs (p=2.25e-12), but only marginally significant for LMOs (p=0.046). Sample quality, as measured by detection p-value, showed strong effects on both frequent HMOs and LMOs, however LMOs were much more influenced (p=8.09e-30) than HMOs (p=3.58e-8). Moreover, the first genetic principal component became a significant factor (p=7.65e-5) when analyzing frequent HMOs, while it had no effect on LMOs (p=0.92) (Table 2).

**Figure 2.**
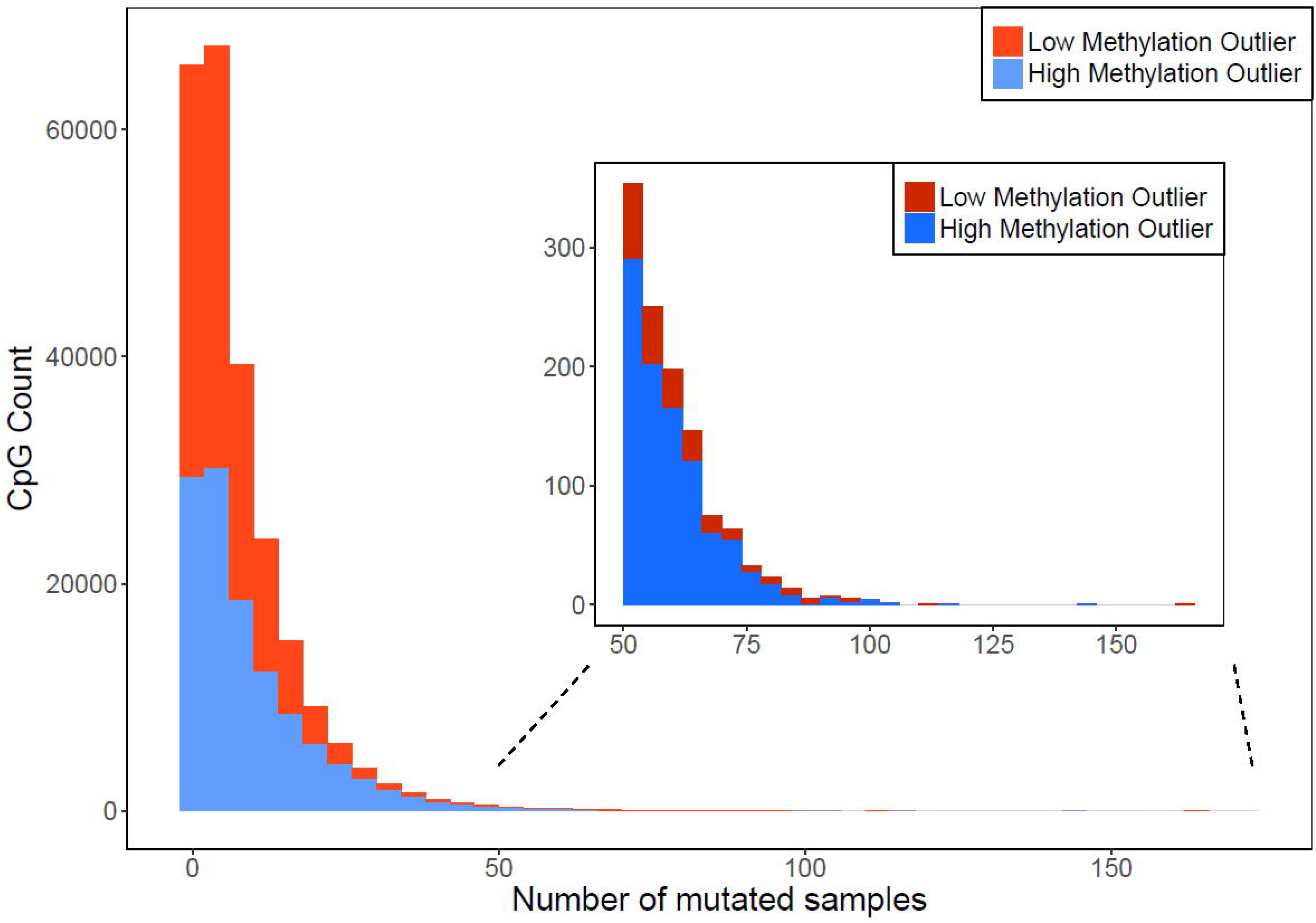
The distribution of mutated samples for high methylation outliers (HMOs) and low methylation outliers (LMOs). For most CpGs, epigenetic mutations only occurred in a small number of samples, but HMOs were more likely to appear in a large number of samples (n>50) than LMOs (969 HMOs and 216 LMOs, p<1e-16).

### Functional annotation of epigenetic mutations

To characterize HMO and LMOs, we examined their locations in relation to CpG island regions and regulatory features. Compared to all CpGs analyzed, where 33.5% of CpGs locate in CpG islands, HMOs were enriched within CpG islands (63% of CpGs, p<1e-16) and frequent HMOs even more so (88% of CpGs, p<1e-16). On the other hand, LMOs were mostly located outside of CpG islands (88% CpGs outside of CpG islands, p<1e-16), but the opposite was true for frequent LMOs, which were enriched in CpG islands (51% of CpGs, p=8.6e-8) (Figure 3). We further explored regulatory features of the frequent epigenetic mutations using the Ensembl database [10], and found that frequent HMOs were enriched in promoter regions (p=1.1e-10), but less likely to be found in CCCTC-Binding factor (CTCF) binding sites (p=1.4e-09) and regions of open chromatin (p=3.6e-07) (Figure 4A). The frequent LMOs, on the other hand, were enriched in CTCF (p=7.7e-12) and transcription factor binding sites (p=3.9e-05), open chromatin (p=0.0012), and promoter flanking regions (p=0.041), while depleted in promoter regions (p=6.9e-19) (Figure 4B).

**Figure 3.**
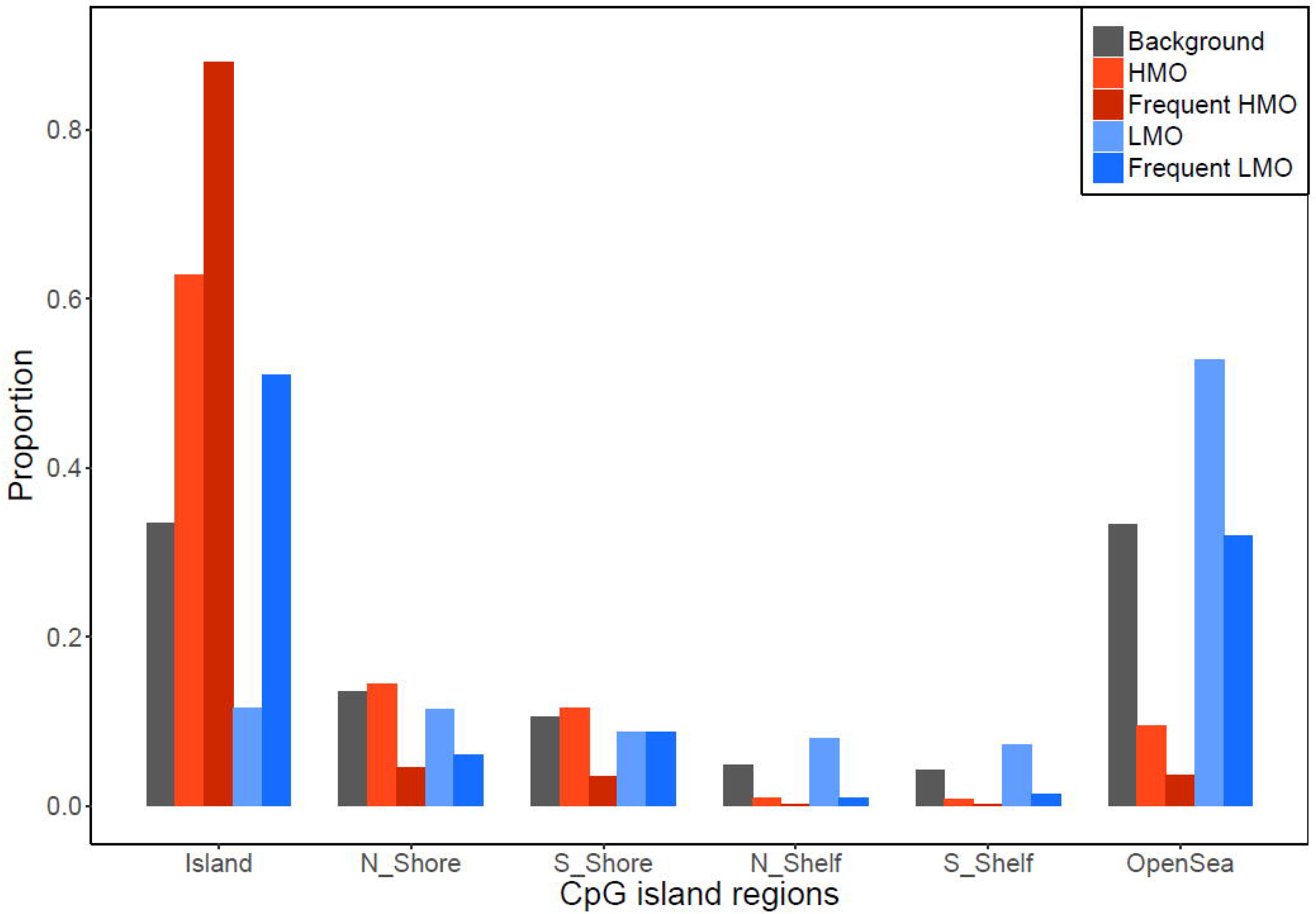
Proportions of high methylation outliers (HMOs) and low methylation outliers (LMOs) in different CpG island regions. HMOs are enriched in CpG islands (p<1e-16) while LMOs are more distributed outside of CpG islands (p<1e-16), especially in open sea regions. However, both frequent HMOs and LMOs are enriched in CpG islands (p<1e-16 and p=8.6e-8).

**Figure 4.**
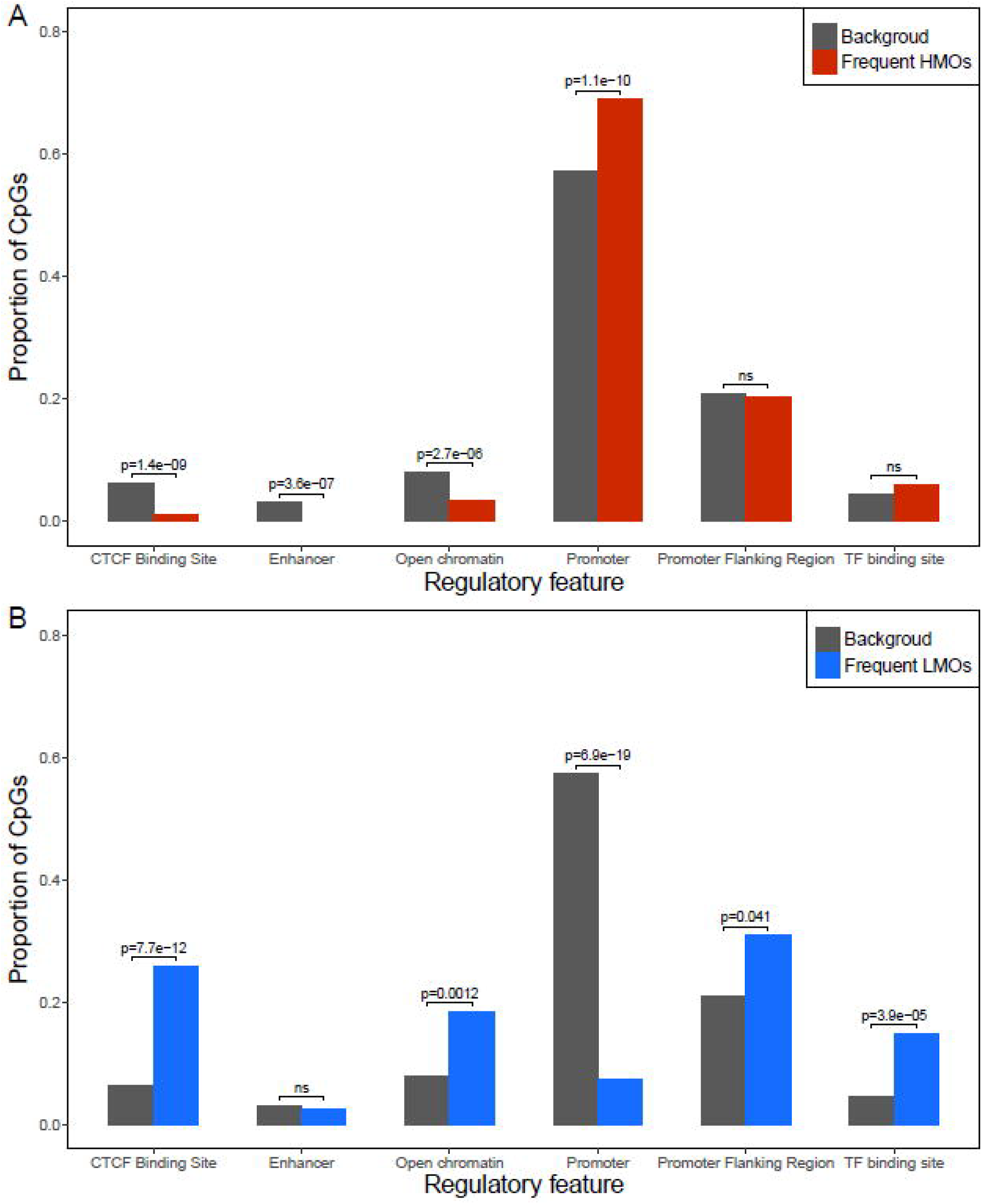
The distribution of regulatory features of frequent high methylation outliers (HMOs) and low methylation outliers (LMOs). Compared to the background distribution of the 450k array design, frequent HMOs were enriched in promoter regions (A), while the opposite was true for LMOs (B).

### Epigenetic mutation is associated with cancer diagnosis

As aberrant DNA methylation levels in gene regulatory regions may cause abnormal gene expression, which may be associated with cancer, we analyzed epigenetic mutations in relation to cancer diagnosis in the SATSA participants. Cancer diagnosis date was retrieved using linkage to The National Patient Registry (prior to May 2016) including ICD-codes for all cancer types (ICD7 codes 140-205, ICD8 codes 140-209, ICD9 codes 140-208, ICD10 codes C00-C97 and B21). The SATSA participants included 29 prevalent cancer cases diagnosed already at baseline, and 79 incident cases that developed cancer during the follow-up period. Hence, information on whether the participant was diagnosed with cancer by the end of the follow-up was tested in the mixed model for associations with log10-transformed numbers of epigenetic mutations. Samples of individuals with cancer, including samples before and after cancer diagnosis, were observed to have a higher number of frequent HMOs (p=0.013), but no associations were found for frequent LMOs (p=0.71, Table 2). Furthermore, in the survival analysis, people with a higher number of frequent HMOs had a higher risk of cancer incidence (Table S1).

### Epigenetic mutations are shared within twin pairs

By applying a co-twin control design we could further study the genetic effect and the genetic-age interaction in association with epigenetic mutations. We calculated the number of shared epigenetic mutations within a twin pair sampled at the same time, and studied their association with time and twin zygosity using a random effects model (Table 3). The numbers of shared epigenetic mutations were normalized in order to compare the effect sizes from different sets of CpGs. First, taking all CpGs into account (n=390,894), the number of shared epigenetic mutations increased significantly with age (*β*=0.019, p=0.026) and MZ pairs shared more epigenetic mutations than DZ pairs (*β*=1.078, p=3.41e-18). After excluding 20,660 cis-meQTL CpGs, the age effect became stronger (*β*=0.025, p=5.98e-3) while the zygosity effect was smaller (*β*=0.855, p=1.05e-11). Last, within the 20,660 cis-meQTL-CpGs, the number of shared epigenetic mutations was not associated with age (*β*=2.86e-4, p=0.969), while the zygosity difference (*β*=1.461, p=8.34e-28) was larger than in results from non-meQTL-CpGs. None of the three tests showed significant twin zygosity-age interaction or sex effect.

**Table 3.** The results of the scaled number of shared epigenetic mutations calculated from different sets of CpGs in association with age, sex, twin zygosity and zygosity-age interaction.

|  | Covariates | Estimate | Standard Error | P-value |
| --- | --- | --- | --- | --- |
| All CpGs<br>(390,894) | Age | 0.019 | 8.59e-3 | 0.026 |
|  | Sex | 0.208 | 0.107 | 0.055 |
|  | Zygosity (DZ) | -1.078 | 0.105 | 3.41e-18 |
|  | Zygosity (DZ)×Age | -0.012 | 0.011 | 0.284 |
| Non-cis-meQTL<br>CpGs (370,234) | Age | 0.025 | 9.17e-3 | 5.98e-03 |
|  | Sex | 0.183 | 0.116 | 0.117 |
|  | Zygosity (DZ) | -0.855 | 0.114 | 1.05e-11 |
|  | Zygosity (DZ)×Age | -0.013 | 0.012 | 0.263 |
| Cis-meQTL CpGs<br>(20,660) | Age | 2.86e-4 | 7.61e-3 | 0.969 |
|  | Sex | 0.194 | 0.107 | 0.071 |
|  | Zygosity (DZ) | -1.461 | 1.105 | 8.34e-28 |
|  | Zygosity (DZ)×Age | -3.77e-3 | 9.64e-3 | 0.696 |

meQTL: methylation quantitative trait loci

### Epigenetic mutations were validated using pyrosequencing

To verify epigenetic mutations identified from 450k array, we selected four frequently mutated CpGs (One HMO: cg05270750, and three LMOs: cg17338133, cg25351353, cg05124918) in 93 samples from 26 individuals for validation with pyrosequencing. In general, the pyrosequencing results were well correlated with methylation data measured by the 450k array (cg05270750: r=0.84; cg17338133: r=0.59; cg25351353: r=0.80; cg05124918: r=0.77). In addition, we compared methylation levels of mutated samples to the normal group using results from the 450k array and pyrosequencing respectively. In pyrosequencing data, significant differences were observed between mutated samples and normal ones, using the same definition of a mutated sample as that for the 450k array data (Table 4). Hence, pyrosequencing technically validated epigenetic mutations identified from the 450k array. Although the agreement between the two methods was generally good, we still observed large differences between pyrosequencing and 450k data in some samples, where four samples in cg17338133 and six samples in cg 05124918 showed over 15% methylation level differences between 450k array and pyrosequencing data after centering their mean methylation levels. This indicates that we might wrongly-detect or fail to detect epigenetic mutations from 450k chip data. In general, pyrosequencing data were more stable and changes in methylation levels were smoother than that from 450k array (Figure 5). For example, in cg05270750 measured by the 450k array (Figure 5E), one participant was identified to have epigenetic mutations in the first three measures, but the methylation level turned back to normal status in the last two measures. However, pyrosequencing data showed the methylation levels of the five measures from this individual were consistently defined as epigenetic mutations.

**Table 4.** Results from t-tests comparing methylation levels in samples with epigenetic mutations to normal samples using data from the 450k array and pyrosequencing.

| Data | Number of samples |  | Mean difference<br>(Methylation level,<br>%) | p-value |
| --- | --- | --- | --- | --- |
|  | Normal | Mutation |  |  |
| cg05270750, 450k-chip | 81 | 12 | 13.39 | 4.34e-6 |
| cg05270750, Pyroseq |  |  | 10.79 | 2.01e-3 |
| cg17338133, 450k-chip | 76 | 17 | 13.11 | 6.39e-8 |
| cg17338133, Pyroseq |  |  | 9.35 | 0.02 |
| cg25351353, 450k-chip | 67 | 26 | 14.58 | 7.93e-17 |
| cg25351353, Pyroseq |  |  | 12.70 | 9.20e-8 |
| cg05124918, 450k-chip | 63 | 30 | 21.87 | 3.22e-20 |
| cg05124918, Pyroseq |  |  | 11.08 | 3.76e-07 |

**Figure 5.**
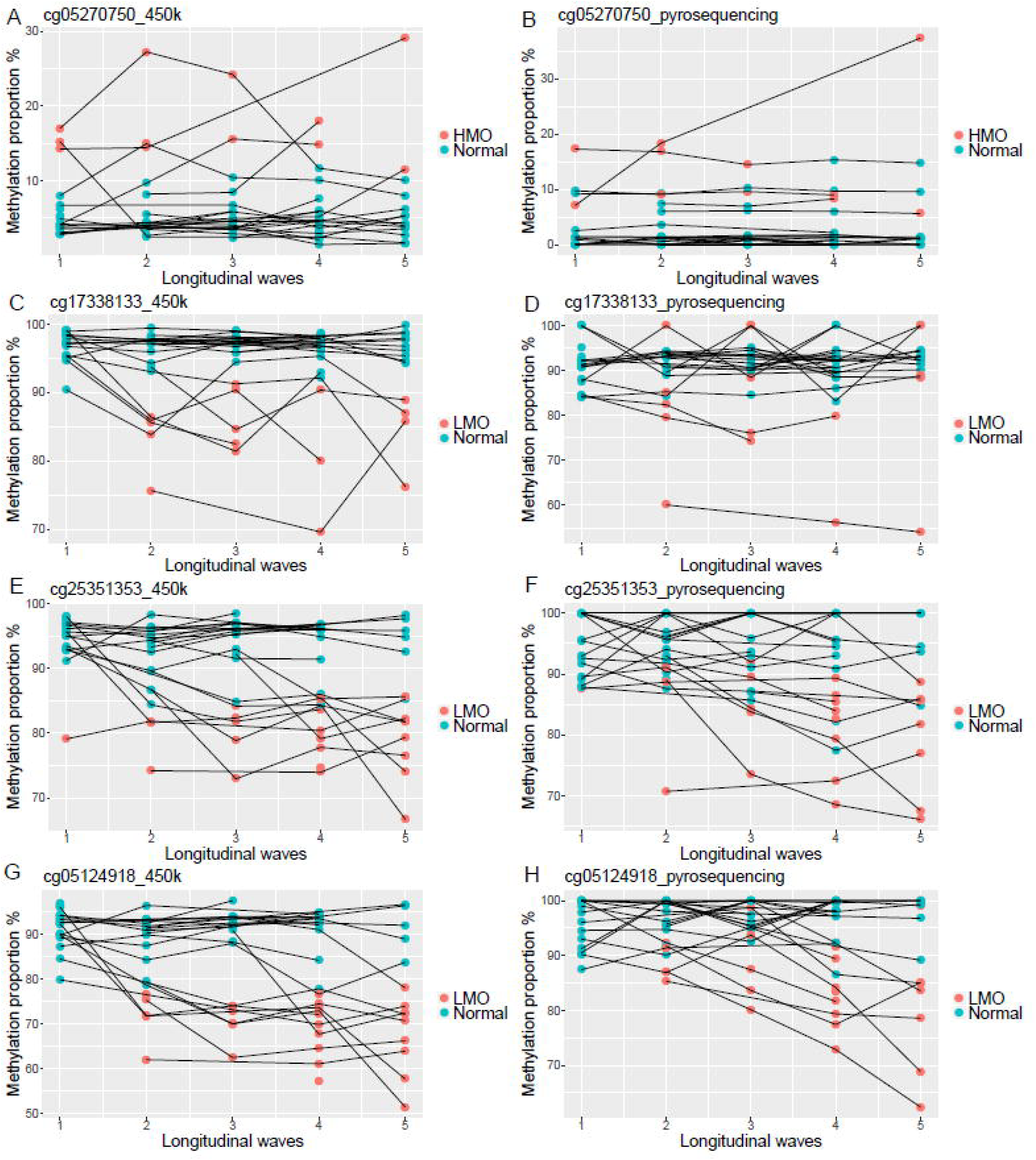
The longitudinal change of four CpGs in 93 samples from 26 individuals measured by 450k array (left panel) and pyrosequencing (Pyroseq, right panel) techniques. Methylation levels of A) cg05270750 from 450k-chip, B) cg05270750 from Pyroseq, C) cg17338133 from 450k-chip, D) cg17338133 from Pyroseq, E) cg25351353 from 450k-chip, F) cg25351353 from Pyroseq, G) cg05124918 from 450-chip, H) cg05124918 from Pyroseq. Samples are shown as points colored by their mutation status defined by the 450k data and lines links longitudinal samples collected in the same individual.

### Functional validation of epigenetic mutations in cancer tissues

To further verify the overabundance of epigenetic mutations in cancer tissues, we picked a gene PR/SET domain 7 (*PRDM7*) which was the only gene related to CpGs tested in pyrosequencing (cg05270750), and analyzed DNA methylation and gene expression data of the gene in tumor tissues and normal adjacent tissues using The Cancer Genome Atlas (TCGA) [11] data downloaded from Wanderer [12]. We selected the four most common cancer types in both sexes combined: lung cancer, breast cancer, colorectal cancer and prostate cancer [13]. The total numbers of tumor and normal adjacent samples were 2,209 and 261 respectively, all cancer types combined. On average, the expression levels of *PRDM7* were higher in tumor tissues than normal adjacent tissues in all cancer types, but the difference was only statistically significant for lung cancer (p=1.83e-09, Table S2). For DNA methylation data, the tumor tissues had significantly lower methylation levels than normal adjacent tissues in the gene body (Figure 6A). However, for CpGs in the *PRDM7* promoter (from cg06295223 to cg26935333), there was no significant difference between the mean methylation levels of cancer and normal adjacent tissues (Figure 6A). To quantify and compare epigenetic mutations in both tissues, we used the distribution of normal adjacent samples to determine epigenetic mutation cutoffs. By calculating the number of epigenetic mutations in tissue samples, tumor tissues had higher proportions of epigenetic mutations in the gene body, while epigenetic mutations were not observed in normal adjacent tissues. In the gene promoter, tumor and normal adjacent tissues had similar and relatively low proportions of epigenetic mutations (Figure 6B).

**Figure 6.**
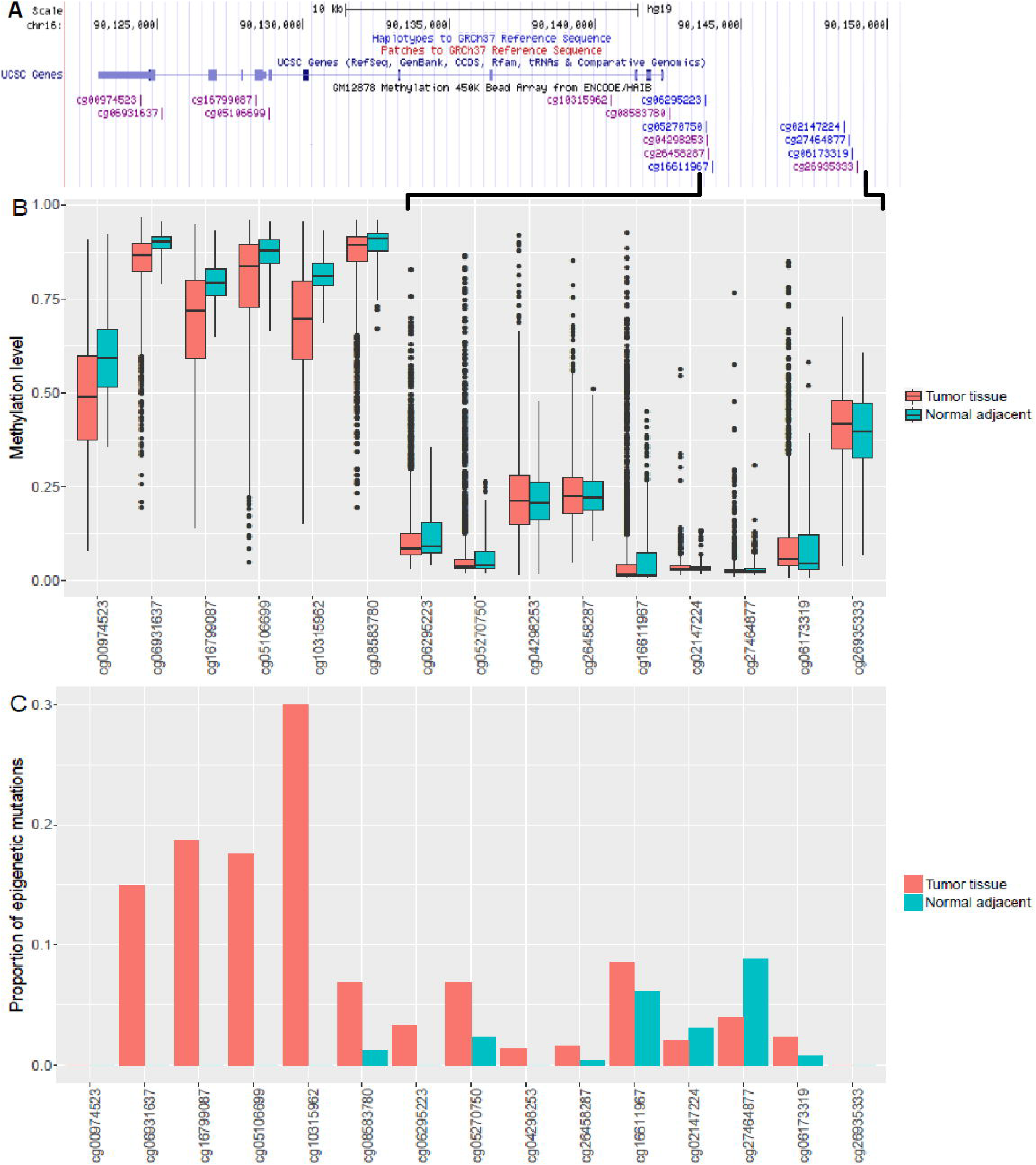
Comparing the DNA methylation and epigenetic mutation patterns of gene *PRDM7* between tumor and normal adjacent tissues. Data were downloaded from TCGA through Wanderer. The cancer types included lung cancer, breast cancer, colorectal cancer and prostate cancer. A) The location of CpGs related to gene *PRDM7* in UCSC genome browser. B) The methylation levels of CpGs in gene *PRDM7*. Tumor and normal adjacent tissues had similar methylation levels in the gene promoter, while the methylation levels of tumor tissues in the gene body were significantly lower than normal adjacent tissues. C) The proportion of epigenetic mutations in tumor and normal adjacent tissues. Tumor tissues had higher proportions of epigenetic mutations in the gene body, while both tumor and normal adjacent tissues had similar but low proportion of epigenetic mutations in the gene promoter.

## Discussion

In this study, we analyzed age-related accumulation of epigenetic mutations from a longitudinal perspective in old Swedish twins. Apart from being exponentially associated with age, epigenetic mutations were also associated with sex, CD19+ B cell count, genetic background, cancer incidence and technical factors. We further analyzed frequent HMOs and LMOs separately and found that biological factors, including B cell compositions and genetic factors, were more strongly associated with frequent HMOs than LMOs, while LMOs were more influenced by technical factors. Moreover, cancer diagnosis was significantly associated with the increase of epigenetic mutations, especially among frequent HMOs, while the same was not true for LMOs. Emerging evidence indicate that epigenetic mutations could be related to cancer [6], as epigenetic mutations may cause abnormal gene expression, which could contribute to the development of cancer. On the other hand, mutated DNA sequences and abnormal epigenetic regulation in tumor cells may in turn cause more epigenetic mutations. In this study, we found that the number of epigenetic mutations was significantly higher in samples of individuals who were diagnosed with cancer by the end of follow-up. Therefore, we conclude that the number of epigenetic mutations may accumulate long before the diagnosis of cancer. The survival analysis further showed that a higher number of frequent HMOs could be a risk factor for cancer incidence. These results support a previous finding where the number of epigenetic mutations were higher in tumor tissues than in normal tissues [8]. Follow-up studies with more participants are needed to better establish the possible relationship between epigenetic mutations and cancer.

In this study, DNA methylation data were corrected for cellular compositions predicted by the Houseman method [14], yet imputed CD19+ B cell count was significantly associated with epigenetic mutations, but not other cell types. A possible explanation could be that B cells have a unique methylation pattern compared to other lymphocytes [15]. Also, B cell composition was still a strong factor for frequent HMOs but the effect became very week for frequent LMOs, probably because cell specific CpGs are enriched in promoter regions [15] where HMOs are mostly found.

When studying functional annotations associated with the epigenetic mutations, we found that the location and regulatory features were different for frequent HMOs and LMOs. The observed enrichment of HMOs in CpG islands and promoter regions indicated that HMOs were more related to biological function than LMOs, which is in line with the fact that technical bias was significant in LMOs.

The concept of epigenetic mutations should be discussed in relation to methylation variability, as they both describe methylation divergence between individuals. However, epigenetic mutations refer to more extreme methylation levels carried by a small number of individuals, while methylation variability is considered to be a population pattern. In contrast to traditional association studies on methylation levels, where CpGs of higher variances are more likely to have statistical power, CpGs of high variances could have too large inter quartile ranges to be identified as epigenetic mutations by definition. Therefore, the identified frequent epigenetic mutations were different from the age-associated CpGs or age-varied CpGs reported prior to this study using the same data [3,5], and thus may contribute to the aging processes by other ways than through the epigenetic drift.

Even after excluding cis-meQTL CpGs, a small genetic effect captured by the first genetic PC was associated with epigenetic mutations, especially in frequent HMOs. To further explore how genetic background and age affected the accumulation of epigenetic mutations, we studied the number of shared epigenetic mutations between twins over time. Here we did not simply exclude cis-meQTL CpGs, but considered them as epigenetic mutations caused by genetic variants inherited at birth. For all CpGs and non-meQTL CpGs, we observed both age and genetic effect associated with the number of shared epigenetic mutations within the twin pair. To isolate the genetic effect, we specifically analyzed cis-meQTL CpGs and found that in this selection, the number of shared epigenetic mutations did not change with age. This result was consistent with a previous paper showing that meQTL-CpG associations are stable over time [16]. Additionally, we failed to detect an interaction between genetic factors and age, indicating that the increase of epigenetic mutations with age was not dependent on the genetic background. Therefore, the remaining genetic effect observed after removing cis-meQTL CpGs was probably due to trans-meQTLs or unidentified cis-meQTLs. In conclusion, the age effect on the accumulation of epigenetic mutations is independent of genetic background. However, we might not have enough statistical power to detect a significant age-genetic interaction on shared epigenetic mutations, since the age effect estimated for MZ twins was larger than for DZ twins. Moreover, due to the limit of the age range in this study (48 to 98 years), we could not exclude the possibility of genetic-associated development of epigenetic mutations in early ages, which remains to be examined by future studies.

Technical artifacts and poor sample quality could lead to erroneous measures that interfere with identifying true biological methylation outliers. Although sample quality control based on detection p-value was applied in the pre-processing pipe-line of the methylation data, it was still found to strongly influence the identification of the epigenetic mutations. Although the technical effect was strong and hard to avoid, the effect of age on epigenetic mutations was not biased as we randomized samples on microarrays. Another important technical artifact is the batch effect from different arrays, but we adjust for batches both in data pre-processing and as a random effect in the mixed effect model. Hence, despite the confounding issues from different technical biases when analyzing methylation outliers, the underlying biological phenomenon of increasing number of epigenetic mutations with age still holds.

Validation of the epigenetic mutations identified in 450k data was done by pyrosequencing, which also detected aberrant methylation levels proving that they were true biological outliers and not simply technical errors. However, some samples showed very different results between the two methods suggesting measurement errors existed. When comparing results from the two methods, pyrosequencing data were more stable and better indicated that epigenetic mutations were persistent over time, which supported the accumulation of epigenetic mutations as a factor of aging.

The HMO site cg05270750 validated by pyrosequencing is located in the promoter region of the gene *PRDM7*, which encodes a Histone-Lysine Trimethyltransferase involved in histone modification. To further explore the potential consequence of epigenetic mutations, we analyzed DNA methylation and gene expression of gene *PRDM7* in data on tumor and normal adjacent tissues from TCGA. The expression of PDM7 in normal adjacent tissues were very low, as previously seen [17]. Nevertheless, we observed higher expression of *PRDM7* in tumor tissues, especially in lung cancers, suggesting the abnormal expression of *PRDM7* could be related to the dysregulation of histone modification in tumor. On the other hand, we observed similar proportions of epigenetic mutations between tumor and normal adjacent tissues in the gene promoter, but more epigenetic mutations in the gene body for tumor tissues. Since normal adjacent tissue can be regarded as an intermediate state between healthy and tumor tissues, it is suggested that, in the process of cancer development, epigenetic mutations were likely to first accumulate in gene promoters and then spread to the whole epigenome.

## Conclusions

In summary, using longitudinal DNA methylation data, we showed that the accumulation of epigenetic mutations is exponentially associated with age in old adults, and once mutations are established, they are stable over time. Furthermore, epigenetic mutations are enriched in important regulatory sites, e.g. promoter regions of genes involved in histone modification processes, which could potentially be an explanation to why people who develop cancer have more epigenetic mutations than others do. In addition, we showed that the burden of accumulation associated with the human aging process is unlikely to be driven by underlying genetic background. Hence, accumulation of epigenetic mutations is an underexplored area in the field of aging, and warrants further studies to enhance our understanding of this phenomenon.

## Methods

### Study population

Twins as participants in this study were enrolled in the SATSA longitudinal cohort study [18]. After quality control, a total of 994 blood samples obtained from 375 individuals in five longitudinal waves (1992-2012) were used in the analyses. The 375 participants had a mean age of 68.9 years (SD=9.7) at their first measurement, and 223 (59.5%) were women. Of the 375 participants, 197 contributed samples in three or more waves. Phenotype data were collected through comprehensive questionnaires and physical testing at each sampling wave. Phenotypes used in this study include chronological age, sex, zygosity, smoking status and cancer diagnosis.

### DNA methylation data

DNA methylation data were obtained from DNA extracted from whole blood samples measured by Infinium HumanMethylation450 BeadChips. In total 485,512 CpG sites were measured for each sample. The quality control and preprocessing methods of the methylation data were described in a previous study [3]. Samples from individuals lacking genetic data were removed, retaining a total of 994 samples for analyses. Blood cellular compositions were estimated by the Houseman method [14] using a reference panel [15]. The methylation data were adjusted by cellular compositions using a linear regression before the analyses. Additionally, batch effects, which were detected as slides on the 450k chip, were adjusted using the Combat method from the sva package [19].

### Genotype data and imputation

Genetic data were measured by Infinium PsychArray (Illumina Inc., San Diego, CA, USA) with 588,454 SNPs detected for every individual. After quality control, data were imputed to the 1000 Genomes Project phase 1 version 3 reference [20] using IMPUTE2 version 2.3.2 [21,22] with default parameters. The first 10 PCs were calculated based on a linkage disequilibrium pruned set of directly genotyped autosomal SNPs.

### Identifying epigenetic mutations

The definition of an epigenetic mutation was consistent with Gentilini *et al* [7]. For each CpG, the quartiles of methylation levels were calculated for every CpG using the first observation available from each individual, and were calculated separately for men and women to avoid the sex effect on methylation levels. Samples having methylation levels three times the inter quartile range higher than the third quartile or lower than the first quartile were identified as mutated outliers. Methylation levels were presented in beta-values, which indicate the methylation proportions. CpGs associated with cis-meQTLs (<1 Mbps) were removed from further epigenetic mutation analyses. For the rest of the CpGs, outlier samples were identified as epigenetic mutations, and the total number of epigenetic mutations was counted for every sample. Identified epigenetic mutations were classified into HMOs and LMOs according to whether they exceed the upper or lower boundary of normal methylation levels (defined as 3 times IQR higher than the third quantile or lower than the first quantile).

### Statistical analysis

A mixed effect model was fitted to measure the association of the number of epigenetic mutations on age and other factors (Equation 1). A log-10 transformation was applied to the number of epigenetic mutations to form a distribution closer to a normal distribution. For each sample, the log10-transformed number of CpGs with detection p-values over 0.01 was used to indicate the sample quality. In the formula, i, j and k denote individual, slide batch and waves; β0, β1, β2, β3, β4, β5, β6 denote fixed intercepts, fixed coefficient of age, sex, CD19 B cell composition, first genetic principal component, detection p-value and whether the individual developed cancer; u0, u1 and ε denotes random intercept of individual, slide batch and random error.

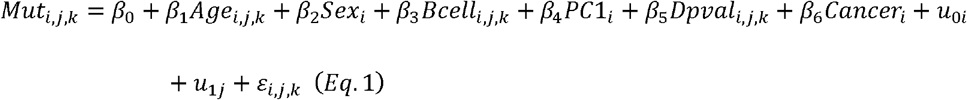

The survival analysis of cancer diagnosis and epigenetic mutations was performed using a Cox model. The model included sex, current smoking as baseline exposure, number of epigenetic mutations as a time-varying covariate, and attained age as the time scale. The model was further adjusted for twin pair and batch effect using robust standard error.

In twin analysis, a mixed effect model was used to study the number of exact same epigenetic mutations between paired twins measured at the same time in association with age, sex and twin zygosity (Equation 2),

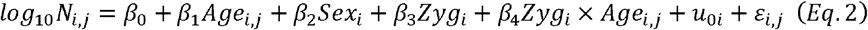

where *i* and *j* denote individual and longitudinal measure; β0, β1, β2, β3, β4 denote fixed intercept, fixed coefficient of age, sex, zygosity and zygosity-age interaction; *u*_0i_, and *ε* denote random intercept of individual and random error.

All statistical analyses were performed in R version 3.4.3.

### Pyrosequencing

In total, 93 samples from 26 individuals were measured by pyrosequencing to validate epigenetic mutations in 4 CpGs (cg05270750, cg17338133, cg25351353, cg05124918). The samples were selected to present 4 to 5 longitudinal measures for every individual. The selection of CpGs was based on their primer quality, and having large numbers of mutated samples. The primers of the four CpGs were designed using the software PyroMark Assay Design by QIAGEN. DNA samples were converted by bisulfite reaction performed on EZ-96 DNA Methylation-Gold™ MagPrep kit provided by ZYMO RESEARCH CORP. Converted samples were randomized in a 96-well plate and sequenced for each CpG on PyroMark Q96 ID provided by QIAGEN. The raw data were processed in PyroMark Q24 Software v2.5.8 by QIAGEN.

## Declarations

### Availability of data and material

The datasets generated and analyzed during the current study are available in Array Express database of EMBL-EBL (www.ebi.ac.uk/arrayexpress) under accession number E-MTAB-7309.

### Authors’ contributions

SH, NP and YW conceived and designed this study. YW performed data processing, statistical analysis and drafted the manuscript. YW and MA conducted pyrosequencing for validation. SH, ÅH, RK, JJ, IK and MA contributed to the manuscript writing. All authors read and approved the final manuscript.

### Ethics approval and consent to participate

All participants in SATSA have provided written informed consents. This study was approved by the ethics committee at Karolinska Institutet with Dnr 2015/1729-31/5.

