## Supplementary files for "Comprehensive longitudinal study of epigenetic mutations in aging"

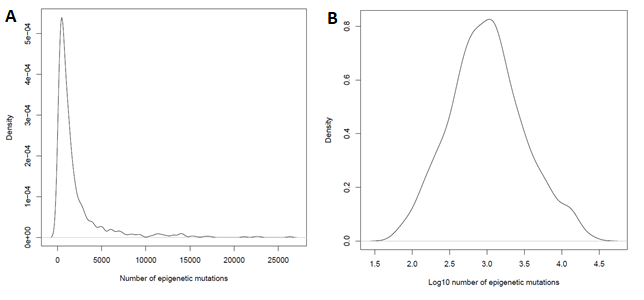

**Figure S1**. The distribution of number of epigenetic mutation. A) The number of epigenetic mutations showed a right-skewed distribution. B) After log-transformation, the distribution is close to normal distribution.

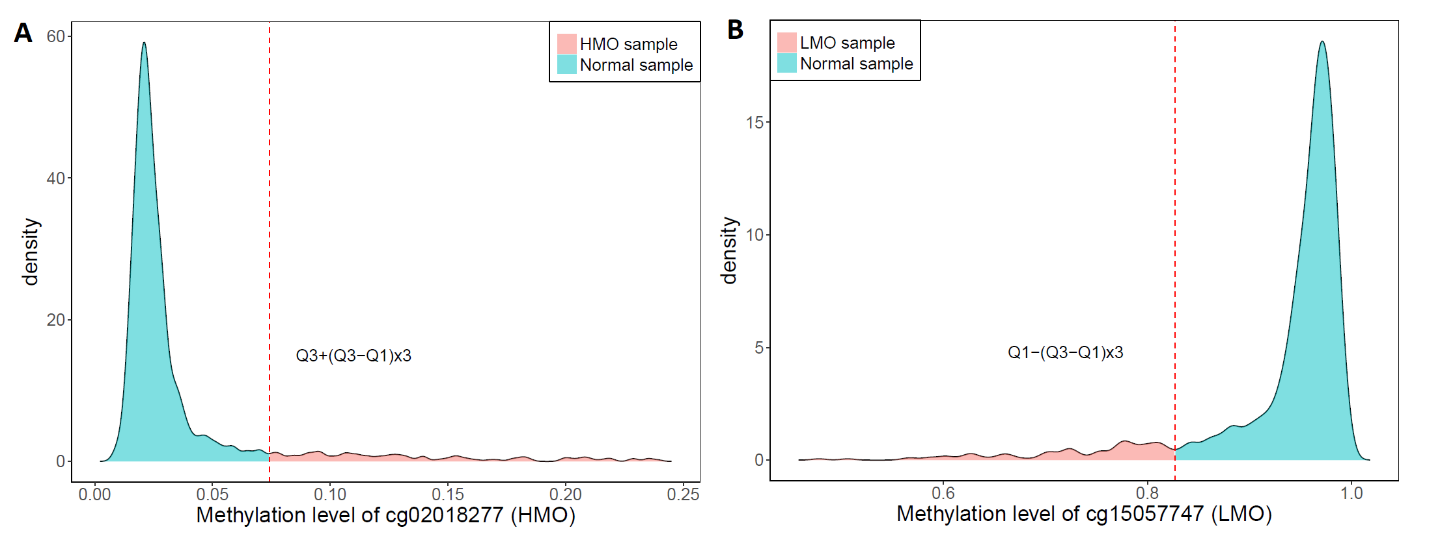

**Figure S2.** The examples of epigenetic mutations in two directions. The distribution of methylation levels of A) cg02018277 (higily methylated outlier) and B) cg15057747 (lowly methylation outlier). The methylation levels of epigenetic mutations were greatly different to normal samples, exceeding three times of inter quantile range of the first or third quantile. The samples in red had epigenetic mutations and their methylation levels greatly differed to normal samples in blue.

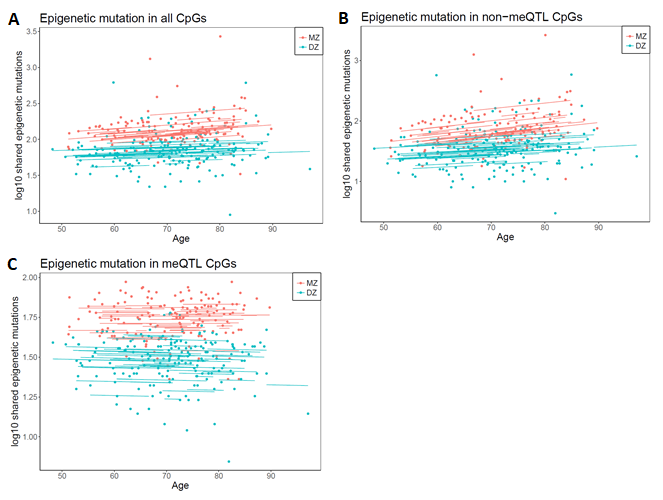

**Figure S3**. The number of epigenetic mutations shared between twins in association with age and twin zygosity. The number of shared epigenetic mutations significantly increased with age for A) all CpGs and B) non-meQTL CpGs. In both case, monozygotic twins (MZ) shared more epigenetic mutations than dizygotic twins (DZ), indicating genetic effect. C) But for meQTL CpGs, epigenetic mutations shared by twins was not associated with age, which suggested that meQTLs were stable over time. None of the three regressions showed age-zygosity interaction, indicating that genetic does not influence the rate of age-associated increase of epigenetic mutations.

**Table S1.** The survival analysis of epigenetic mutations in association with cancer incidence using a Cox proportional hazard model. For epigenetic mutations, frequent HMOs and frequent LMOs the hazard ratios represent the ratio of an increase of 10 epigenetic mutations.

|  | Covariate | Hazard ratio | p-value |
| --- | --- | --- | --- |
| All epigenetic mutations | Female sex | 0.609 | 0.037 |
|  | Current smoker | 1.945 | 0.027 |
|  | Epigenetic mutations | 1.000 | 0.203 |
| Frequent HMOs | Female Sex | 0.623 | 0.044 |
|  | Current smoker | 1.931 | 0.029 |
|  | Frequent HMOs | 1.046 | 0.048 |
| Frequent LMOs | Female Sex | 0.613 | 0.040 |
|  | Current smoker | 1.966 | 0.025 |
|  | Frequent LMOs | 0.939 | 0.757 |

**Table S2.** The mean expression levels of gene *PRDM7* in tumor and normal adjacent tissues for different cancer types. A t-test was used to compare the mean expression levels in tumor and normal adjacent tissues. The gene expression data were downloaded from TCGA through Wanderer.

| Cancer type | Mean expression level (Number of samples) | | P-value |
| --- | --- | --- | --- |
|  | Tumor tissue | Normal adjacent tissue |  |
| Breast invasive carcinoma | 0.237 (720) | 0.205 (84) | 0.439 |
| Colon adenocarcinoma | 0.114 (249) | 0.086 (19) | 0.577 |
| Lung adenocarcinoma | 0.336 (422) | 0.000 (21) | <1e-16 |
| Lung squamous cell carcinoma | 0.323 (359) | 0.179 (8) | 0.245 |
| Prostate adenocarcinoma | 0.227 (333) | 0.122 (34) | 0.063 |
| Combined | 0.255 (2083) | 0.147 (166) | 3.08e-5 |
